# LarvaTagger: Manual and automatic tagging of *Drosophila* larval behaviour

**DOI:** 10.1101/2024.03.18.585197

**Authors:** François Laurent, Alexandre Blanc, Lilly May, Lautaro Gándara, Benjamin T. Cocanougher, Benjamin M.W. Jones, Peter Hague, Chloé Barré, Christian L. Vestergaard, Justin Crocker, Marta Zlatic, Tihana Jovanic, Jean-Baptiste Masson

**Author notes:** Equal contributions (François Laurent, Alexandre Blanc).

## Abstract

**Motivation:** As more behavioural assays are carried out in large-scale experiments on *Drosophila* larvae, the definitions of the archetypal actions of a larva are regularly refined. In addition, video recording and tracking technologies constantly evolve. Consequently, automatic tagging tools for *Drosophila* larval behaviour must be retrained to learn new representations from new data. However, existing tools cannot transfer knowledge from large amounts of previously accumulated data. We introduce LarvaTagger, a piece of software that combines a pre-trained deep neural network, providing a continuous latent representation of larva actions for stereotypical behaviour identification, with a graphical user interface to manually tag the behaviour and train new automatic taggers with the updated ground truth.

**Results:** We reproduced results from an automatic tagger with high accuracy, and we demonstrated that pre-training on large databases accelerates the training of a new tagger, achieving similar prediction accuracy using less data.

**Availability:** All the code is free and open source. Docker images are also available. See git-lab.pasteur.fr/nyx/LarvaTagger.jl.

## I. INTRODUCTION

*Drosophila* larva is increasingly used as an animal model for large-scale experiments in behavioural neuroscience [6, 10, 13, 19]. Such assays require automating multiple processing steps, from preprocessing the video streams to generating a behavioural readout suitable for comparing experimental conditions and interpreting the observed differences.

Characterising the behaviour of highly deformable blob-shaped animals such as larvae has been challenging, in contrast to animals that exhibit distinguishable limbs, extensions or body parts. For example, deep learning techniques have been notably successful in identifying behavioural patterns, sometimes even without expert annotation, by tracking the distinct parts of structured-bodied animals [11].

In large-scale behavioural experiments, a camera grabs the arena from above and shows a dorsal view of several dozen of *Drosophila* larvae. In most cases, their body segments are not clearly distinguishable. The limited information on the shape of individual larvae and the large amount of video data has motivated a multi-stage approach to behaviour extraction. Tracking procedures are applied first, often in the online regime, so only dynamic contours and spines are saved without the original video stream. Behaviour characterisation is a separate offline processing step, where a time series of body postures is used as input.

Behaviour is typically characterised by such time series with an implicit function in mind: moving forward, turning, withdrawing, etc. Movement amplitude and speed are useful features to distinguish between actions. However, if the presumed function of the behaviour is given priority over the continuous representation of the motion, an expert may identify the same behaviour at different space and time scales.

Different experimental protocols will rely on the use of *Drosophila melanogaster* strains with different genetic backgrounds and/or various genetically modified lines that can exhibit different behavioral characteristics with similar behavioral categories displaying various dynamics, amplitudes, duration, and sometimes sequences. In addition, environmental context, like the crawling surface or temperature, will also influence larval behavior. It is thus necessary to be able to classify behavioural actions taking into account these different types of behavioural variation.

For example, a hunch, a type of defensive action that larvae perform typically in response to mechanical stimuli, by retracting their head, is characterised by a faster head retraction than the otherwise similar movement of the head and thoracic segments during a peristaltic crawl. Time-dependent features would naturally be selected to distinguish a hunch from a specific phase of the peristaltic wave. However, some genetically modified larvae are slower in both types of movements, so their hunches are slower than other larvae’s peristaltic waves. Consequently, in this latter case, the successful identification of a hunch would preferably rely on duration-invariant features.

Generally speaking, each experimental paradigm introduces different constraints on characterising behaviour. This makes the case for a deep-learning approach to action identification, with discriminating features automatically extracted from the raw time series of postures on a per-experiment basis.

We introduce LarvaTagger, a software tool for action identification based on a pre-trained neural network that can be retrained on new data and actions. LarvaTagger also features a graphical user interface to visualise tracking data and manually tag actions. The first section of this paper briefly lists key features and use cases for LarvaTagger. We discuss its contribution to the software ecosystem and the technologies it is based on. In the second section, we characterise the performance of a tagger trained to reproduce the behavioural readout of a widely-used tagger [10], with similar changes in action probabilities for most larva lines. We also demonstrate transfer learning by first pre-training a neural network on a database in a self-supervised fashion and, second, training for a particular tagging task on another database.

Throughout the present article, we also make a general argument in favour of continuous representations of the observed behaviour as a complementary approach to the discrete behavioural readout given in terms of actions. The conclusion stresses how LarvaTagger can facilitate the combination of both approaches.

## II. SOFTWARE ELEMENTS AND METHODS

LarvaTagger is a behaviour tagging tool for *Drosophila* larvae. It performs both manual and automatic tagging, with the possibility to train new taggers using a pre-trained neural network and labelled/tagged “ground-truth” data.

The LarvaTagger project is divided into multiple sub-projects to support different use cases. For example, the user interface (UI) is provided by LarvaTagger.jl, available at gitlab.pasteur.fr/nyx/larvatagger.jl. The LarvaTagger.jl repository is the main entry point of the whole project and its documentation. However, the automatic tagging logic is functionally independent from the UI. We expect this to make the design of alternative taggers easier. In the present article, LarvaTagger is demonstrated in combination with a tagger based on an autoencoder known as MaggotUBA [2].

For compatible platforms^1^, LarvaTagger is preferably downloaded and run as a Docker image from Docker Hub (hub.docker.com/repository/docker/flaur/larvatagger). Past versions are available to ensure a high degree of reproducibility. Standalone scripts for Windows (cmd and PowerShell), macOS and Linux help operate the Docker image. Figure 1a identifies other pieces of software LarvaTagger can be combined with or that it can replace. In particular, the original motivation for LarvaTagger was to replace Pipeline_action_analysis_t5_pasteur_janelia (referred to in the following as [Pipeline_pasteur_janelia](https://github.com/DecBayComp/Pipeline_action_analysis_t5_pasteur_janelia), [10]), a tagger that identifies specific actions but cannot be adjusted to new data or new actions.

**FIG. 1.**
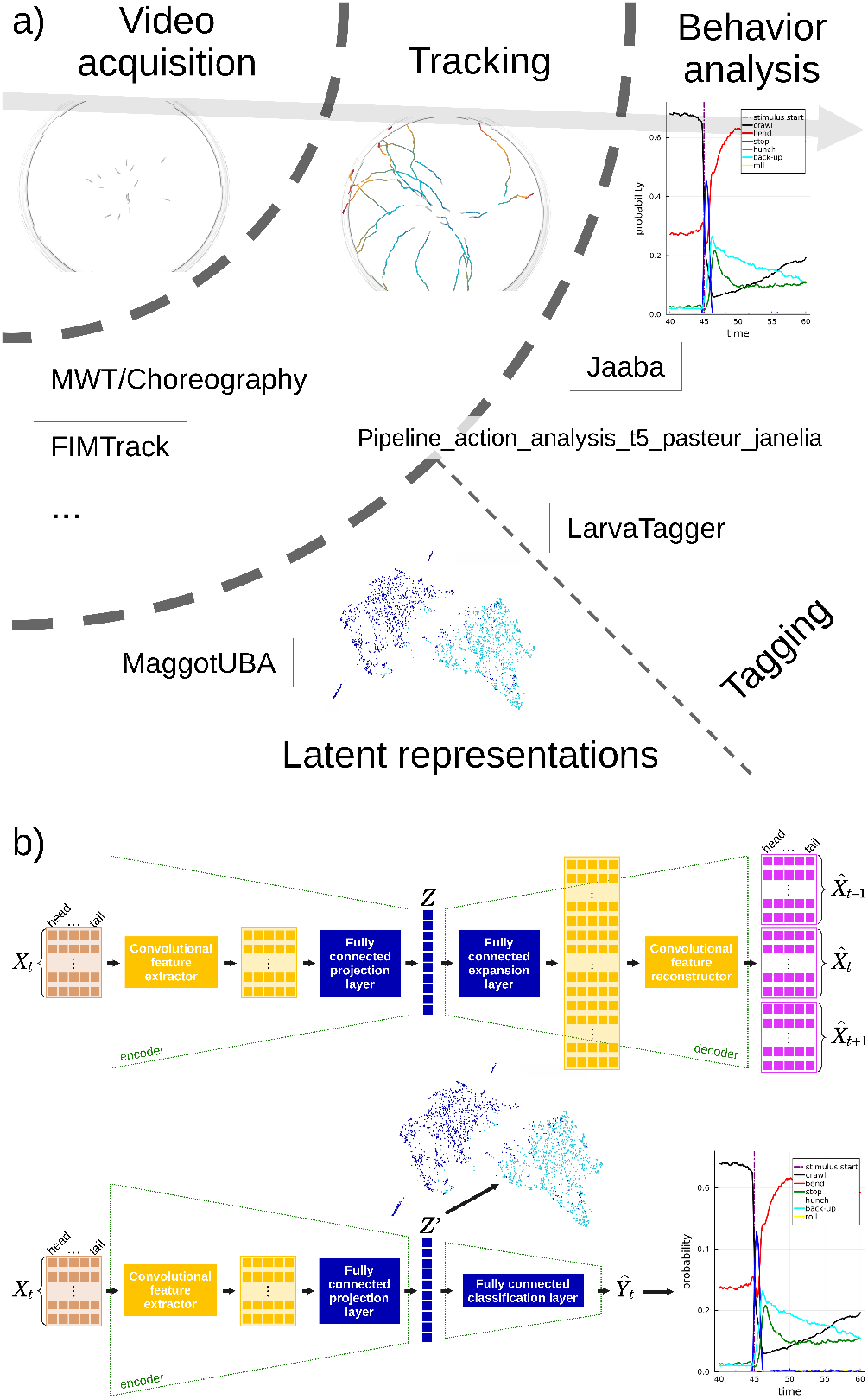
a) Tools and approaches to go from video data to behaviour representations: the moving objects in video data (an example frame is shown in the top left corner) are tracked with tools such as MWT or FIMTrack (see text for references). Behaviour analysis is carried out using tracking data (top, middle; colours show time), with tools such as Jaaba, [Pipeline_pasteur_janelia](https://github.com/DecBayComp/Pipeline_action_analysis_t5_pasteur_janelia) or LarvaTagger. Here, [Pipeline_pasteur_janelia](https://github.com/DecBayComp/Pipeline_action_analysis_t5_pasteur_janelia) is identified primarily as a tagging tool but also includes a head-tail reorientation procedure pertaining to the tracking domain. Behaviour is analyzed at the population level, typically in discrete actions (*e*.*g*. count or frequency data like in the top right illustration). Alternatively, behaviour can be analyzed using continuous representations in latent spaces. Sequences of postures are projected in a low dimensional latent space as depicted in the bottom illustration of a cloud of 2D points (one point = one sequence) with the colour encoding different actions. MaggotUBA is specifically designed for this approach, and LarvaTagger’s automatic tagging backend is based on MaggotUBA. b) Integration of MaggotUBA (top diagram), an autoencoder that extracts features of the continuous behaviour in a self-supervised fashion. A sequence of postures *X*_*t*_ is compressed into a low-dimensional latent representation *Z* from which a longer sequence, including past and future postures in addition to the input *X*_*t*_, is reconstructed. Training the autoencoder allows learning behavioural features that compress the dynamics in the latent space. The encoder features are reused in combination with a classification stage in the MaggotUBA-based tagger (bottom diagram) to learn a discrete behavioural dictionary. The MaggotUBA-based tagger can be used to generate both types of behavioural readouts.

### A. Tracking

LarvaTagger supports several data formats of tracking data, including spine/outline files from MWT/Choreography [17], table.csv files from FIMTrack v2 [16] and MatLab files from [Pipeline_pasteur_janelia](https://github.com/DecBayComp/Pipeline_action_analysis_t5_pasteur_janelia) [10]. File converters have also been written for HDF5 files from Tierpsy Tracker ([5]; see also github.com/Tierpsy/tierpsy-tracker) and DeepLabCut [8].

Tracking tools must be suitable for *Drosophila* larvae and should generate not only point tracks (as illustrated at the top of Fig. 1a) but also posture data, in particular the larval mid-line for automatic tagging purposes, and the contour for visualisation and manual tagging purposes.

Tracking *Drosophila* larvae is challenging because of the lack of structure in the 2D shape of a larva when seen from above. Furthermore, the choice of the number of larvae per behavioural assay represents a trade-off between increasing statistics and minimising crossing events. Indeed, the larvae have highly deformable bodies and are challenging to properly differentiate and track when they interact while being filmed at low resolutions (recent progress have been made at high resolution [18]). All the tracking solutions we know of require precise calibration and may fail to detect moving objects with minor variations of experimental parameters.

While larva tracking is out of LarvaTagger’s scope, we leveraged mwt-container, an automatic tracking pipeline for the batch processing of video data. mwt-container is conveniently available as a Docker image, easily convertible into a Singularity/Apptainer image file for use on high-performance computing clusters. It takes AVI files as input and applies the Multi-Worm Tracker (specifically mwt-core and [Choreography](https://github.com/Ichoran/choreography), [17]). Unlike other applications, MWT is shipped without its LabView components. Some tracking hyper-parameters are automatically adjusted so that the number of simultaneously tracked objects approaches the number of larvae in the assay. See supplementary section **Tracking** for more details.

### B. Manual tagging

LarvaTagger’s entry point is the LarvaTagger.jl Julia package, available at gitlab.pasteur.fr/nyx/larvatagger.jl, which provides command-line (CLI) and graphical user interfaces (GUI). The GUI is suitable for inspecting and manually tagging tracking data. It features the following capabilities:

- Searching for a track/larva by id.
- Visualising tracking data and any associated label on a per-frame basis or animated at different playback speeds; labels are colour-coded and the contour of the tracked larva is coloured accordingly.
- Assigning labels for individual larvae at each defined time step or across entire time segments.
- Defining new labels, renaming labels, changing the associated colour, *etc*.
- Undoing all manual editions performed on a larva.
- Editing the metadata of the behavioural assay in a dedicated panel.
- Exporting the labelling/tagging information of selected tracks; the desired tracks can be selected from another panel indicating which tracks have been manually edited.

The labelling/tagging information is stored in a JSON file whose structure is very similar to the *Worm tracker Commons Object Notation* (WCON) format [5] for tracking data, with specifications available at gitlab.pasteur.fr/nyx/planarlarvaefiles. JSON files are human-readable, suitable for storing metadata in addition to data, and easy to load and edit in every programming language, or even using a text editor.

### C. Automatic tagging

LarvaTagger can automatically tag the behaviour in discrete actions, either from the GUI for the currently loaded data file or in batch mode using the CLI. LarvaTagger separates the UI and general tagging API from the core logic of the tagger. This separation should make it easier to implement other tagging backends. By default in the Docker images, LarvaTagger ships with a tagging backend based on MaggotUBA, called MaggotUBA-adapter. In turn, this latter backend ships with a default tagger, which currently is called `20230311` and is demonstrated in section III below.

The tagging backend supports a default tagger and can train new taggers with different target actions and ground-truth data. Conceptually, a tagger is an instance of the model implemented by the backend. Training a new tagger can only be performed using the CLI or the Python API of MaggotUBA-adapter.

The LarvaTagger project is divided into multiple sub-projects; some of these are listed below:

- LarvaTagger.jl: command-line and graphical user interfaces, which is the recommended entry point.
- TaggingBackends: backbone for operating and designing tagging backends; manages data preparation and trained taggers.
- MaggotUBA-adapter: implements a MaggotUBA-based tagger as illustrated in 1b; can be operated directly as a library in Python.
- PlanarLarvae.jl: low-level logic for handling data files and sampling in data repositories; specifies the JSON file format used to store tagging information and metadata.

A map of these sub-projects is found in the developer documentation available at gitlab.pasteur.fr/nyx/larvatagger.jl/-/blob/dev/doc/develop.md.

Figure 1a also highlights a similar tool with manual tagging and automatic tagger retraining: Jaaba [7]. However, a key distinction of LarvaTagger from Jaaba is in its intermediate latent representations of the behaviour, which does not necessarily depend on data annotations or discrete actions as defined by behaviour experts. Indeed, as illustrated in Fig. 1b, LarvaTagger relies on MaggotUBA ([2], implemented with PyTorch [15]), a self-supervised autoencoder that can project short sequences of postures into a common low-dimensional latent space. As showcased in [2], the latent representations can be used to implement a statistical test, or can be inspected in regions of the latent space where higher between-group variance is observed.

LarvaTagger trains new taggers using these latent representations to feed a classifier with automatically extracted features. The latent representations can evolve if the training procedure is allowed to fine-tune the encoder neural network with the purpose of action classification tasks (Fig. 1b). In addition to identifying discrete actions, a MaggotUBA-based tagger can generate these latent representations for the input data. This can be performed using the CLI.

## III. EMULATING AN EXISTING TAGGER

As mentioned above, a motivation for LarvaTagger was to replace [Pipeline_pasteur_janelia](https://github.com/DecBayComp/Pipeline_action_analysis_t5_pasteur_janelia) [10], a tagger that identifies specific actions. Indeed, this latter tagger cannot be retrained because it was incrementally designed following an active learning approach: predictive components were added to the tagger as new needs arose, *e*.*g*. new actions were to be identified, or corrections to be performed [3, 9]. In addition, each component takes as input not only explicit features of the data, but also predictions from the preexisting components, thus forming a hierarchical ensemble of classifiers. While the hierarchical design has helped enforce priorities of some actions over others (in particular to handle some actions so rare as to be only identified in some controlled circumstances or genetic lines of *Drosophila*), adapting the tagger to new data or new actions is challenging, and may require substantial redesign.

[Pipeline_pasteur_janelia](https://github.com/DecBayComp/Pipeline_action_analysis_t5_pasteur_janelia) identifies 6 actions [10]: forward crawl (*crawl* ), backward crawl (*back* or *back-up*), head cast or bend (*bend* ), hunch, roll and stop. Each action can be further labelled as small or large (or weak or strong). The small/weak actions are grouped under a single *small action* class, resulting in a total of 7 classes.

### A. Reproducing qualitative results

In [10], the authors found neurons of interest by inactivating them in an assay where they subjected larvae to an the air puff as mechanical stimulus [6]. They identified these neurons by observing statistically significant differences in both the frequency of actions and transitions between actions in response to sensory stimulation. This comparison was made between larvae with inactivated neurons and reference larvae (*w;;attP2* ). In total, 293 genetic lines were found to induce significant behavioural variations. These data come from a screening experiment in [6, 10].

We trained a tagger (referred to as `20230311`) based on MaggotUBA on data from the same optogenetic screen used to train the reference tagger in [10]. We analyzed the same selected lines from [6, 10] and mimicked the evaluation procedure of [10], selecting a 1-s time window right after the stimulus onset at 45 s, and applying a similar statistical procedure (*χ*^2^ tests, Bonferroni-corrected for 471 comparisons), to characterise the increases or decreases in population probabilities for each action. The training and evaluation procedure is further detailed in supplementary sections **The 20230311 tagger** and **Evaluation on [6**, **10]**.

Figure 2 illustrates side-by-side results from each tagger. For example, figs. 2a and 2b show that the proportions of the different actions are well preserved in control larvae, especially before the stimulus onset. In response to the air puff, the overall change in proportions (or probabilities) is larger with the MaggotUBA-based tagger than with the original tagger. Figures 2c and 2d show how some *Drosophila* lines compare with the control line, in terms of increase or decrease in action probabilities, and how the MaggotUBA-based tagger reproduces most of these differences.

**FIG. 2.**
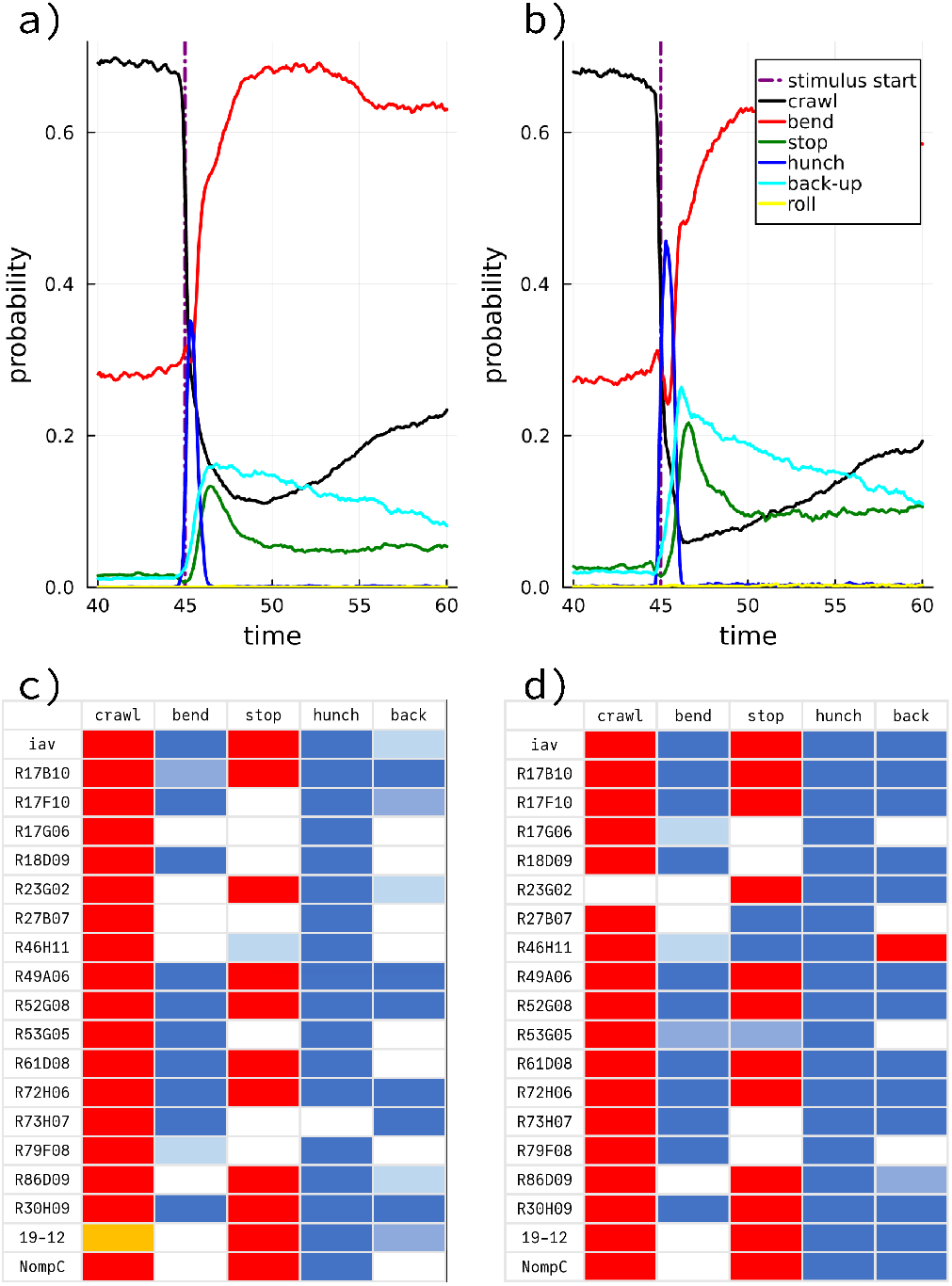
Left panels originate from [Pipeline_pasteur_janelia](https://github.com/DecBayComp/Pipeline_action_analysis_t5_pasteur_janelia). Right panels originate from `20230311`. a-b) Probability time series for the 6 actions of interest (*small action* not accounted for) in a population of control larvae (*w;;attP2* ). At *t* = 45*s*, the larvae received a 38-s long air puff, resulting in a dramatic change in behaviour reflected in the action probabilities over time. While baseline probabilities are well preserved in b), as compared with a), short-term response probabilities exhibit slightly more overall changes in b), with more frequent hunches, back-ups and stops. c-d) Significant differences in individual action probabilities between a selection of *Drosophila* lines and the control *w;;attP2* line. Red denotes a significant increase with *p <* 0.001 and orange an increase with *p <* 0.05. The darkest shade of blue denotes a significant decrease with *p <* 0.001, the intermediate shade of blue a decrease with *p <* 0.01 and the lightest blue a decrease with *p <* 0.05. All *p*-values are Bonferroni-corrected for 471 comparisons. c) is a reproduction of fig. 3e in [10]. Some differences between c) and d) can be observed. In particular, a few effects in c) are lost in d), and, more frequently, effects observed with the 20230311 tagger (d) were not found using the original tagger (c).

Of the 293 comparisons shown in the figs. 3e, 3f, 3g and 5a of [10], we reproduced 232 (79%), considering a three-level outcome for the individual comparisons: positive difference, negative difference, or absence of effect. In most of the remaining cases, the MaggotUBA-based tagger led to the identification of an effect (significant difference) while the original tagger did not (218 significant differences in total *versus* 168, respectively). The only clear discordance (opposite effect) we found involves line *R27B12* : the proportion of *back* was expected to be significantly higher in that line than in control larvae, as reported in [10] fig. 5a, while the MaggotUBA-based tagging led to observing a lower proportion.

**FIG. 3.**
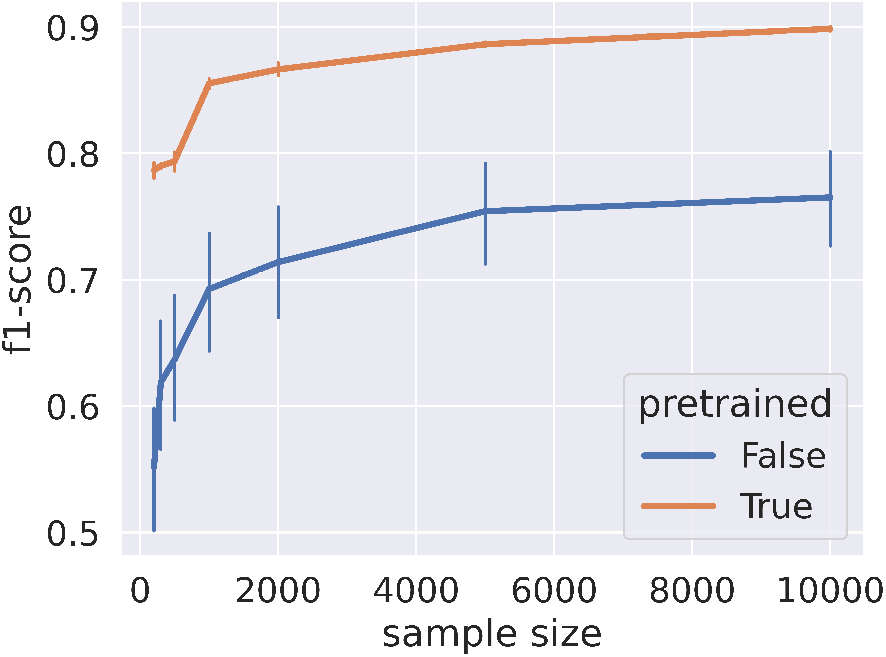
*f* 1-scores for different training dataset sizes (abscissa), with the MaggotUBA encoder pre-trained (orange) or not (blue). Train and test datasets were drawn from the new optogenetic screen. Pre-training was performed on [6, 10].

**FIG. 4.**
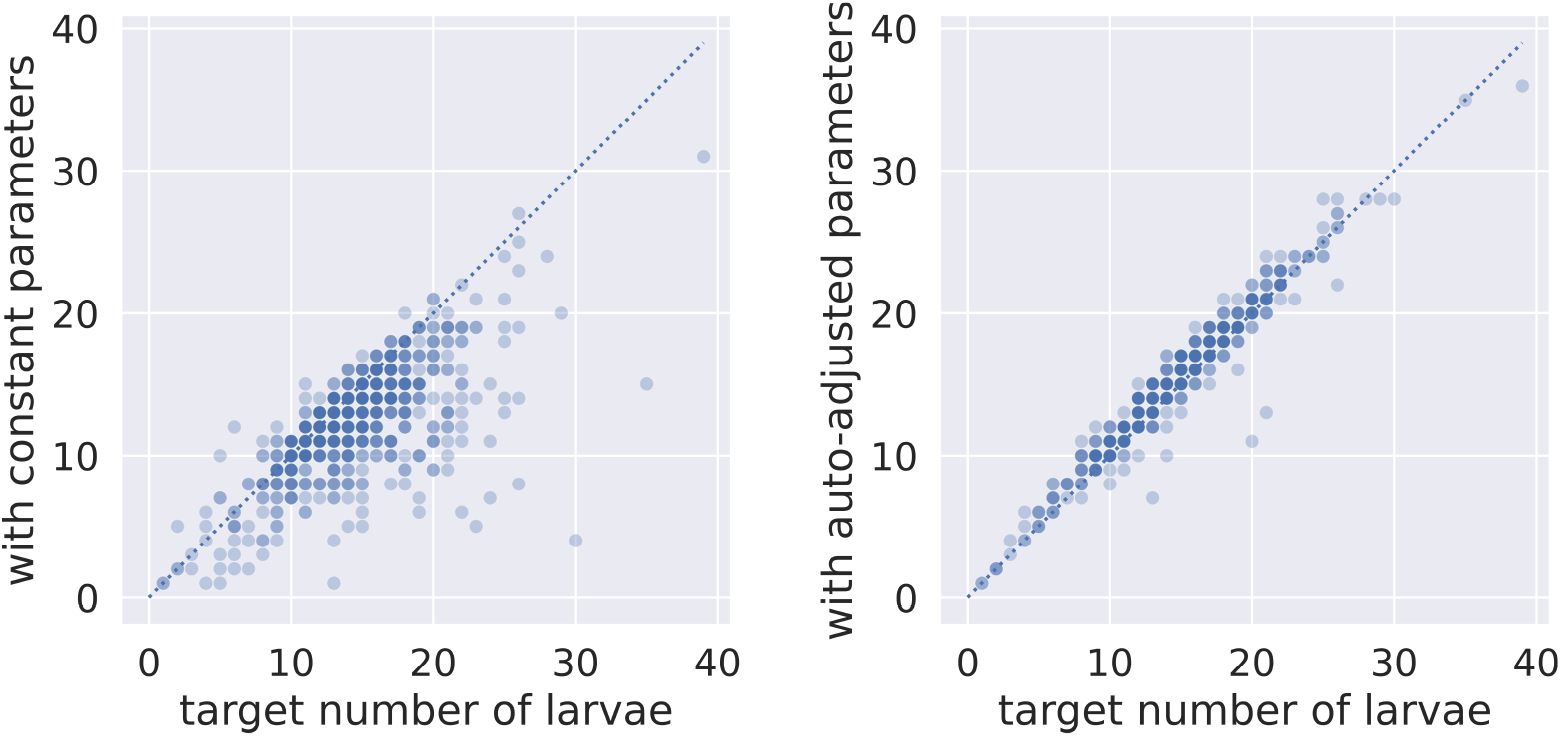
Maximum number of simultaneously tracked larvae (ordinates) in every assay among 699 video files versus the known number of larvae (abscissa). Each transparent blue dot represents one or more assays. Ideally, they should align along the *x* = *y* axis.

All the actions described so far (included in fig. 2) are confirmed (*large*/*strong* ) actions. Regarding the undecided (*small* /*weak* ) actions, we observed that the MaggotUBA-based tagger was more perplexed overall, predicting more such small actions than the original tagger. For example, in the first 1-s window after stimulus onset, these undecided actions represented 29.9% of all the time steps with the MaggotUBA-based tagger and 18.6% with the original tagger.

The differences in behavioural readout between the taggers is most likely explained by the reliance of the MaggotUBA-based tagger on a 2-s time window only, while the original tagger considers entire tracks. In addition, the expert knowledge might be too sparse in the limited training dataset. For example, priority rules introduced in the original tagger made an action be preferably labelled as a *crawl* or *bend*, even with little certainty, rather than as a *hunch* or *roll*.

### B. Transfer learning

The large amount of data accumulated at Janelia Research Campus, such as described in [6, 10, 19], is expected to provide collections of behaviours with unprecedented diversity. Indeed, the different genetic lines of *Drosophila* exhibited differences in their resting behaviour and in their responses to environmental stimuli, hence our interest in taking advantage of the information that the respective data repositories may contain.

The approach consists in pre-training the MaggotUBA autoencoder on these repositories in a self-supervised fashion (no tagging required) to extract general features of the continuous behaviour. Reusing an encoder pre-trained on large amounts of unlabelled data promotes MaggotUBA-based tagger’s ability to achieve higher accuracy when trained with relatively little annotated data (see [1]).

To test this approach, we generated a new dataset in which we activated a randomly chosen set of 50 split-GAL4 lines that express the optogenetic activator of neural activity, Chrimson, in subsets of larval neurons ([12]; see also supplementary section **Optogenetic neural activation screen**). We tested *ca*. 60 animals from each split-GAL4/UAS-Chrimson population. We optogenetically activated neurons in each population for 15 seconds using 660 nm light. Further in this article, we will refer to this dataset as the “new activation screen”, although it actually is a superset of the data used to train the reference tagger in [10].

We pre-trained several MaggotUBA encoders on data from [6, 10], and then trained and tested MaggotUBA-based taggers on small subsets of data from the new optogenetic screen, using the pre-trained encoder to build the MaggotUBA-based taggers. Indeed, data from [6, 10] and the new optogenetic screen feature major dissimilarities that make them interesting candidates to investigate knowledge transferability between distinct experiments. Data from [6, 10] consists of inactivating one or multiple neurons of the larva and characterising the behavioural response to air puffs. On the other hand, data from the new optogenetic screen consists in activating individual or multiple neurons. Consequently, the observed behavioural responses differ in both the nature of actions evoked by the stimuli and their temporal dynamics.

We took the predictions of [Pipeline_pasteur_janelia](https://github.com/DecBayComp/Pipeline_action_analysis_t5_pasteur_janelia) as ground truth. We evaluated transfer learning by comparing the tagging accuracy when using a pre-trained encoder *versus* a naive Xavier-initialised encoder. Further details are given in the **Transfer learning** supplementary section.

Figure 3 shows a dramatic increase in *f* 1-score with a pre-trained encoder at equal training budget. Interestingly, even the smallest datasets help train an initial tagger as long as a pre-trained encoder is used.

Without pre-training, the accuracy exhibits large variations between trials, which may result from a lack of data and insufficient training budget, especially for the larger datasets. Pre-training ensures a more stable training experience and may save time in tweaking the training hyper-parameters. These results are consistent with the literature in self-supervised learning regarding accuracy improvement in downstream tasks with pre-trained neural networks. However, improvements are not systematic and were to be demonstrated in the present application. Indeed, the diversity of the behavioural phenotypes, variability in larval shapes and data quality, and the behavioural drift observed over time could have limited the generalisability of the features learned during pre-training.

In practice, the lower *f* 1-scores in fig. 3 are mainly attributable to *bends* misclassified as other actions and *vice versa*. In particular, most *rolls* were false positives. It is worth noting there are very few *rolls* in the pre-training dataset, according to experts. Still, enough could be found among [Pipeline_pasteur_janelia](https://github.com/DecBayComp/Pipeline_action_analysis_t5_pasteur_janelia)‘s predictions (see inductive bias in [2]). With accumulating user experience from other experiments, training a tagger with a pre-trained encoder may suffer similar defects when the corresponding data exhibit peculiarities such as high tracking noise levels or abnormally slow larvae. In those cases, we found that the training budget should be increased so that the weights in the encoder are fine-tuned to unlearning the patterns seen in pre-training data that do not generalise well to the new training data.

## IV. CONCLUSION

LarvaTagger brings several improvements in the area of *Drosophila* larval behaviour analysis, with a modular design to accommodate various tagging techniques and, at first, approach behaviour in terms of discrete actions.

Currently, it features a pre-trained deep neural network that allows training new taggers with relatively low amounts of data. The same neural network can also generate continuous representations of the behaviour, which opens a new perspective on behaviour characterisation and does not heavily rely on expert annotation.

We argue this is a key advantage because behaviour is a complex concept whose definition is subject to incremental and/or contextual changes. New behaviours can emerge from new experimental paradigms, especially if behaviours are defined in terms of the function they fulfil for the behaving animal (crawling to go forward, bending to change direction, hunching to shelter the sensory centres located in the head, *etc*). More commonly, behaviour is first defined as patterns of movement. However, if man-made, this approach typically results in poorly defined actions, with no specifications of the initiation and termination of the behaviour. Last^2^ but not least, behaviour can also be defined mechanistically, describing how the observed movement patterns are generated. For example, if a neuro-muscular program can be unveiled, behaviour may eventually be decomposed into possibly complex sequences of intermediate actions. Yet, the relationship between neural computation and muscle-mediated behavioural output remains to be properly modelled.

In practice, larval actions and postures have been defined in part with a focus on discriminating between these actions or postures. When asked to annotate the same data examples, the proposed definitions are sometimes too loose for multiple experts to agree on. The difficulty in formalising and discretising behaviour can be circumvented by taking the alternative route of data-driven behaviour analysis, which is paved by approaches such as MaggotUBA. The choice currently depends on the experimental paradigm and modelling goals.

At present, LarvaTagger is being actively used in a study associated with a larval model of Alzheimer’s disease in *Drosophila* larvae (in preparation) and in a large-scale pesticide screening experiment [4].

## Supplementary information

## VI. OPTOGENETIC NEURAL ACTIVATION SCREEN

The optogenetic activation experiment referred to as the “new activation screen” was performed as previously described [6, 14]. For each behavioural assay, *Ca*. 30 larvae were separated from food by bathing them in a 20% sucrose solution for a maximum of 10 minutes. They were rinsed and placed into a square 23 cm^2^ behavior rig covered with 4% agar. We recorded videos of larval behavior, with a DALSA Falcon 4M30 camera for a total of 120 s. At 30 and 75 s, 15 s-long pulses of 660 nm red light (4µW/mm^2^, Philips Lumileds) were applied.

Larvae were tracked in real-time using the Multi-Worm Tracker (MWT) software [13, 17, 19]. We rejected objects that were tracked for less than 5s or moved less than one body length of the larva. For each larva, MWT returns contour and spine coordinates as a function of time. Raw videos are never stored.

## VII. TRACKING

In this section, we briefly describe a tracking pipeline successfully applied to another large-scale screening experiment that required the fully automatic processing of thousands of 1-minute videos, recording freely moving *Drosophila* larvae exposed to different pesticides.

The pipeline is available as a Docker image and can be pulled from quay.io. The image generation code and documentation is available at gitlab.com/larvataggerpipelines/mwt-container.

This pipeline includes:

- mwt-core, a C++ library for online tracking that constitutes the core component of a tracking solution known as Multi-Worm Tracker or MWT [17];
- [Choreography](https://github.com/Ichoran/choreography), a Java utility (part of MWT) for offline processing of the tracks produced by mwt-core;
- some additional C++, Python and Julia code to load video files, using the OpenCV library, and operate mwt-core and [Choreography](https://github.com/Ichoran/choreography) in a head-less fashion (see mwt-cli);
- an optimisation procedure written in Python, using the Optuna library, to automatically adjust the tracking hyper-parameters so that the average number of tracked larvae approaches the target number of larvae (see larva-tagger-tune).

The rationale for adding the optimisation approach is that the number of larvae in the assay is always controlled or measured. If the recording parameters are stable enough, adjusting the tracking parameters may not be required. Otherwise, achieving several tracked objects similar to the known number of moving larvae can guide the automatic selection of the tracking parameters.

Fig.4 shows that in the case constant hyper-parameters do not generalise well to the whole collection of videos, the above-mentioned strategy is an efficient alternative. In the present experiment, we picked 699 videos, manually adjusted the hyper-parameters on 3 of these videos on the one hand (left panel), and automatically adjusted the hyper-parameters on a per-assay basis on the other hand (right panel).

## VIII. THE 20230311 TAGGER

Technical and methodological details of MaggotUBA are provided in [2].

Although LarvaTagger can support substantially different tagging backends, we highlight one particular MaggotUBA-based tagger referred to as `20230311`. `20230311` is composed of a MaggotUBA encoder with a 25-dimension output (*i*.*e*. 25 latent features) and a downstream classifier consisting of a single dense (fully connected) layer. It assigns each data point any of the following 7 labels: *back-up, bend, crawl, hunch, roll, stop* and *small action*. These labels match the terminology used in [10] and — for users of the original tagger — are supposed to match labels from [Pipeline_pasteur_janelia](https://github.com/DecBayComp/Pipeline_action_analysis_t5_pasteur_janelia) as follows:

### 20230311 **labels labels in trx.mat files**

*back-up back large* = *back strong*

*bend cast large* = *cast strong*

*crawl run large* = *run strong*

*hunch hunch large* = *hunch strong*

*roll roll large* = *roll strong*

*stop stop large* = *stop strong small action small motion*

A valid data point is a 2-s time segment of tracking data, can be taken at any defined time step with at least 1 s of past and future history, and consists of a time series of 5-point spines (or mid-lines) re-sampled at 10 Hz around the time step of interest. For prediction tasks, data points in a track’s first or last seconds are labelled in a nearest-neighbour fashion.

As a first step, an autoencoder was trained with self-supervision on a subset of combined data from the new optogenetic screen described in section **Transfer learning** and the screen from [6, 10]. No labelling information was used except for the inductive bias in the pre-training dataset formation that consisted of randomly picking data examples of the different actions so that no classes (or actions) were represented more than twice the least common class. This *×*2 limitation made the pretraining dataset small in comparison with the training dataset, but ensured rare events such as *rolls* were almost equally represented.

In a second step, the resulting pre-trained encoder was combined with the classifier and a new training dataset was sampled from the new optogenetic screen, involving 1 200 235 data examples. These data examples were randomly picked so that no classes were represented more than 20 times the least common class. The combined encoder+classifier was trained with a cross-entropy loss and a 10 000-iteration budget. In the first 5 000 iterations, the classifier only was updated. The encoder and classifier were fine-tuned in the remaining 5 000 iterations.

Similarly to MaggotUBA, training parameters were stored in (2) JSON files and the learned weights in (2) PyTorch PT files.

More details can be found in gitlab.pasteur.fr/nyx/MaggotUBA-adapter#20230311-0-and-20230311.

## IX. EVALUATION ON DATA FROM [6, 10]

The analysis is demonstrated in a code repository available at gitlab.com/larvataggerpipelines/t5_analysis_replicates, in the shape of Pluto notebooks (in Julia).

The same part of the [6, 10] data repository was used as in [10], specifically, lines obtained by crossing with the `UAS TNT 2 0003` effector, and behavioural assays with a 30-s air-puff at *t* = 45 followed by a sequence of 10 2-s air-puffs every 10 s starting from *t* = 105.

A variant of `20230311`, available in MaggotUBA-adapter as `20230311-0`, was applied and compared with [Pipeline_pasteur_janelia](https://github.com/DecBayComp/Pipeline_action_analysis_t5_pasteur_janelia). `20230311-0` is the same tagger as `20230311`; the only difference is a post-processing step in `20230311` that maps the original 12-class output of `20230311-0` (*back strong, back weak, cast strong, cast weak*, etc) onto 7 classes with *weak* actions pooled together in a *small action* class, and some actions renamed (see label table VIII).

The tracking data were extracted from *trx*.*mat* files generated by [Pipeline_pasteur_janelia](https://github.com/DecBayComp/Pipeline_action_analysis_t5_pasteur_janelia), instead of the original files generated by Choreography. Indeed, [Pipeline_pasteur_janelia](https://github.com/DecBayComp/Pipeline_action_analysis_t5_pasteur_janelia) includes a head-tail correction preprocessing step, which technically is a tracking task. This correction step fixed the orientation of roughly 10% of the larvae in the [6, 10] experiment, due to the high numbers of simultaneously behaving larvae and higher probability of U-turn. To better characterise the tagging performance *per se*, we have considered this correction mechanism as common ground for comparing between taggers.

For completeness, running the same analysis on the original data files with head-tail orientation as inferred by MWT+Choreography led to a 10-point increase in baseline level of *back-up* in control larvae, for example. Although the qualitative differences reported in tables figs. 3e, 3f, 3g and 5a in [10] were similarly preserved overall, the higher baseline level of *back-up* probability had no additive effects on known events of higher *back-up* probability. For example, in fig.2b, a 10% plateau can be seen after the stimulus onset, for example, at *t* = 60. The difference in probability between *t* = 40 to *t* = 60 disappeared which — considering the nature of the analyses that can be carried out on such a readout — is a major difference.

## X. TRANSFER LEARNING

Pre-training MaggotUBA autoencoders was performed similarly to `20230311`, here on [6, 10] only, with different latent dimensionalities (25, 50, 100, 200). Training datasets were drawn from the new optogenetic screen, randomly picking 1 000 data files and then a fraction of the available time segments in these files. The corresponding test datasets consisted of similarly sampled time segments from 100 other data files.

The training was performed with a 1 000-iteration budget (default value in MaggotUBA-adapter, subject to changes). The untrained encoders were trained together with the classifier at all iterations. In contrast, with pretrained encoders, the budget was split in two, with 100 iterations to train the classifier initially and 900 iterations to fine-tune the encoder and the classifier jointly.

Fig.3 shows results for a 25-dimension latent space. Very little variation was observed between the explored dimensionalities. An optimum was observed with 50 dimensions.

The source code for transfer learning is available at gitlab.com/larvataggerpipelines/Autoencoding.

## Footnotes

1 At present, an issue seems to affect the Docker images on Apple M2 architecture

2 Evolutionary biology offers a fourth approach to studying behaviour. See Tinbergen’s four questions.

